# Two stages of substrate discrimination dictate selectivity in the *E. coli* MetNI-Q ABC transporter system

**DOI:** 10.1101/2025.01.20.633972

**Authors:** Janet G. Yang, Hulda Yuchun Chen, John H. Guardado, Maile Gardner, Matthew S. Foronda

## Abstract

The *Escherichia coli* MetNI-Q importer, an ATP-binding cassette (ABC) transporter, mediates the uptake of both L- and D-enantiomers of methionine. Original *in vivo* uptake studies show a strong preference for L-Met over D-Met, but the molecular basis of this selectivity is unclear. In this work, we systematically examine substrate discrimination by the MetNI transporter and MetQ substrate binding protein using an array of biophysical and biochemical techniques. Based on the kinetic and thermodynamic parameters of individual intermediates in the transport cycle, we uncover multiple steps in the transport cycle that confer substrate specificity. As in many other ABC importer systems, selectivity is applied at the level of binding to the substrate binding protein: MetQ dictates a 1,000-fold preference for L-Met over D-Met. However, beyond this initial level of selectivity, MetQ displays distinct binding preferences for the MetNI transporter depending on the substrate. We propose that the differences in binding affinities reflect the more favored release of L-Met into the permeation pathway when compared to D-Met. In support of this model, under saturating conditions, MetNI transports L-Met across the lipid bilayer at a faster rate than D-Met. Interestingly, the ATPase activity of the MetNI-Q complex is not modulated by the presence of substrate. Our studies reveal that the MetNI-Q system incorporates two separate steps in tuning methionine uptake to substrate chirality and availability. This method of discrimination ensures the import of the most biologically preferred substrate while also allowing for adaptability to more limiting nutrient conditions.

## Introduction

ABC transporters are found across all kingdoms of life and constitute one of the largest protein superfamilies (1–4). These integral membrane proteins play key roles in crucial cellular functions including transition metal acquisition, nutrient import, cellular detoxification, osmoregulation, membrane organization, and antigen processing (5–9). Using the energy from ATP binding and hydrolysis, they unidirectionally transport a diverse range of substrates against electrochemical gradients. ABC importers, found only in prokaryotes, transport nutrients such as amino acids, sugars, and vitamins (10–13). Exporters, on the other hand, are present in both prokaryotes and eukaryotes and extrude molecules such as xenobiotics, peptides, and lipids (14–17). The study of ABC transporters is highly clinically relevant, as mutations in human ABC transporters have been linked to several conditions, including cystic fibrosis, liver disease, tuberculosis, diabetes, macular degeneration, metabolic diseases, and multi-drug resistance (18–25). Additionally, ABC transporters have been shown to play a role in bacterial and fungal virulence, enabling the uptake of crucial nutrients in organisms such as *Streptococcus pneumoniae*, *Bacillus anthracis*, and *Salmonella enterica* (26, 27).

All ABC transporters contain two highly conserved nucleotide-binding domains (NBDs) located in the cytoplasm and two variable transmembrane domains (TMDs) embedded in the membrane. Progression through the cycle of ATP binding and hydrolysis at the NBDs drives rearrangement of the TMDs to allow for substrate passage across the membrane. Additionally, importers require a cognate substrate-binding protein (SBP) or substrate-binding domains (SBDs) that are found either in the periplasm, anchored to the extracellular side of the membrane, or covalently attached to the TMDs. The canonical model of ABC importer mechanism is shown in Figure 1 (28, 29). In this simplified model, the SBP sequesters the substrate and delivers it to a “resting state” apo-nucleotide transporter. The substrate-loaded SBP and apo-transporter form a stable complex, and subsequent binding of two ATP molecules at the NBDs induces transition of the TMDs to an outward-facing conformation. This rearrangement concurrently drives the release of the substrate into the transmembrane cavity. Lastly, ATP hydrolysis triggers formation of the inward-facing conformation, release of the SBP and transport of the substrate across the membrane.

**Figure 1.**
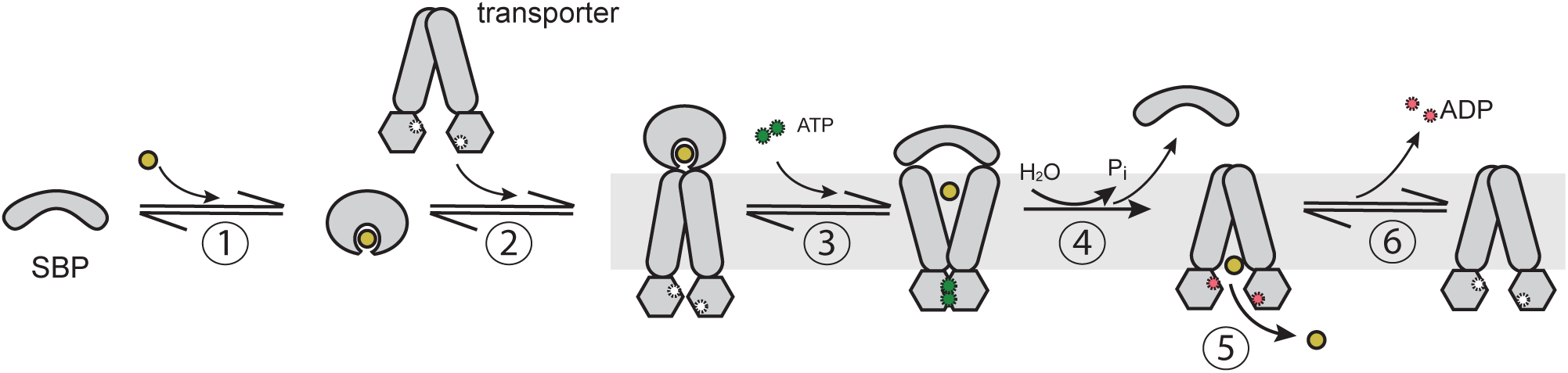
The canonical model of transport. (1) Apo-SBP in the open conformation binds substrate which induces the closed SBP conformation. (2) Substrate-bound SBP docks onto nucleotide-free transporter in an inward-facing resting conformation. The resulting SBP-transporter complex adopts a pre-translocation conformation. (3) The SBP-transporter complex binds ATP, which triggers transporter rearrangement to the outward-facing conformation, concurrent opening of the SBP, and release of the substrate into the transmembrane cavity. (4) ATP hydrolysis drives transition to the inward-facing conformation, thereby stimulating release of the apo-SBP. (5) The transmembrane cavity is now exposed to the interior of the cell, and substrate is released into the cytoplasm. (6) ADP dissociates from the transporter, resetting the system for another round of transport.

While some importer systems are tailored for only one specific substrate, many ABC import systems can transport related substrates as necessary (10). An open question in the field is how these importer systems implement substrate selectivity while also maintaining the flexibility necessary for survival in nutrient-depleted conditions. Several studies proposed that this plasticity is enabled by the cognate SBP or SBDs, many of which can bind more than one substrate (30). SBPs and SBDs have a bilobal architecture connected by a flexible hinge region, and substrate binding prompts closure of the lobes relative to one another via an induced-fit mechanism (31). EPR, smFRET, and MD simulation studies showed that the degree of closure between the lobes can vary based on the available substrate, and thus there are multiple different substrate-bound conformations for some SBPs (30–33). Examples of promiscuous substrate binding include the LivJ SBP, which binds to leucine, isoleucine, and phenylalanine (34). In another amino acid importer system, YecSC-FliY, the SBP captures both L- and D-enantiomers of sulfur-containing amino acids (35). Beyond amino acid transport, the MolA SBP binds both molybdate and tungsten, and the sugar-binding GGBP delivers both D-glucose and D-galactose to the MglABC transporter (36, 37).

While many SBPs are capable of binding numerous substrates, whether or not overall transport selectivity is exclusively dependent on the SBP is unclear. For example, in the Opp oligopeptide transport system, the peptide binding preferences of the OppA SBP matches the overall peptide selectivity of the entire system (38). In contrast, in the maltose import system, the SBP can bind many different maltodextrins with high affinity; however, only a subset can be transported across the membrane (39, 40). Structural studies revealed a well-defined sugar binding site within the TMDs of the maltose transporter that, in combination with several other functional studies, conclusively demonstrated that substrate selectivity is dictated by both the SBP and the maltose transporter (41). If these differing models of selectivity apply more broadly to the field is yet to be determined.

To explore the different possible mechanisms for substrate discrimination, we chose to study the *E. coli* methionine importer system, which is comprised of the MetNI transporter and MetQ SBP (42–45). Elegant *in vivo* studies from almost 50 years ago showed that both L- and D-Met are translocated by this system and that the system is more easily saturable with L-Met (K_m_ = 0.13 µM) compared to D-Met (K_m_ = 1.16 µM) (46). The preference for L-Met is intuitive from a biological viewpoint; the L-enantiomer is ready for use, while the D-form requires isomerization before incorporation into protein synthesis and metabolic pathways. Recent structural and *in vivo* work support a mechanism in which the methionine enantiomers are translocated via two different pathways. In this model, L-Met is transported via the canonical pathway (Fig. 1), while D-Met is transported via a noncanonical pathway. In the noncanonical mechanism, D-Met accesses the permeation pathway of MetNI through a pre-formed MetNI-Q complex (47). Thus far, the kinetic and thermodynamic feasibility of specific models derived from these studies remain largely untested.

Here we use a variety of biochemical and biophysical techniques to systematically dissect the molecular basis of MetNI-Q specificity. We measure the specificity of L- and D-Met binding by MetQ and the effect of the enantiomers on the thermodynamics of MetNI-Q complex stability. We characterize MetNI ATPase activity stimulated by L- and D-Met and directly follow the transport of these substrates using purified components. Furthermore, the kinetic parameters are compared in different lipid environments, including detergent micelles, nanodiscs, and proteoliposomes. Ultimately, we find that L- and D-Met discrimination occurs at two distinct steps in the MetNI-Q transport cycle: substrate binding by the SBP and substrate release induced by formation of a SBP – transporter complex. To the best of our knowledge, the latter represents a new selectivity determining mechanism for an ABC transporter system. Together, these two steps allow *E. coli* to preferentially uptake L-Met even when D-Met is available. However, when L-Met is not present in the environment, the cell can instead rely on MetNI-Q to import the less-preferred D-enantiomer to meet cellular needs.

## Results

### Characterization of MetNI trans-inhibition mutant

In the MetNI-Q transport system, L-Met acts as both a substrate and an allosteric inhibitor (43, 48, 49). In addition to being transported by MetNI-Q, L-Met can bind and allosterically inhibit ATP hydrolysis by binding MetNI at two C2 regulatory domains. This streamlined regulatory mechanism, termed trans-inhibition, limits the uptake of L-Met when intracellular concentrations are adequate for optimal growth. While critical in balancing the import of L-Met, trans-inhibition can complicate the interpretation of functional studies.

To disentangle the two roles of L-Met for our detergent-based and nanodisc studies, we chose to use a previously-identified N295A MetNI mutation that does not bind L-Met at the C2 regulatory domain (47). To ensure that the N295A mutation did not affect other aspects of MetNI function, we compared the ATPase activity of detergent-solubilized wild-type and N295A MetNI as a function of MgATP concentration (Figs. 2 and S1). The kinetic parameters derived from Michaelis-Menten analysis are identical between the two, suggesting that the mutation does not affect ATP usage (Figs. 2a and 2c). Additionally, we measured the inhibitory constants (K_i_) for both L- and D-Met (Fig. 2b). The N295A mutation increased the K_i_ for L-Met by 28-fold and had modest effect on K_i_ for D-Met. Based on this suppressed L-Met inhibition and wild-type ATPase activity, we selected N295A MetNI as a reliable alternative to study MetNI function in the absence of trans-inhibition.

**Figure 2.**
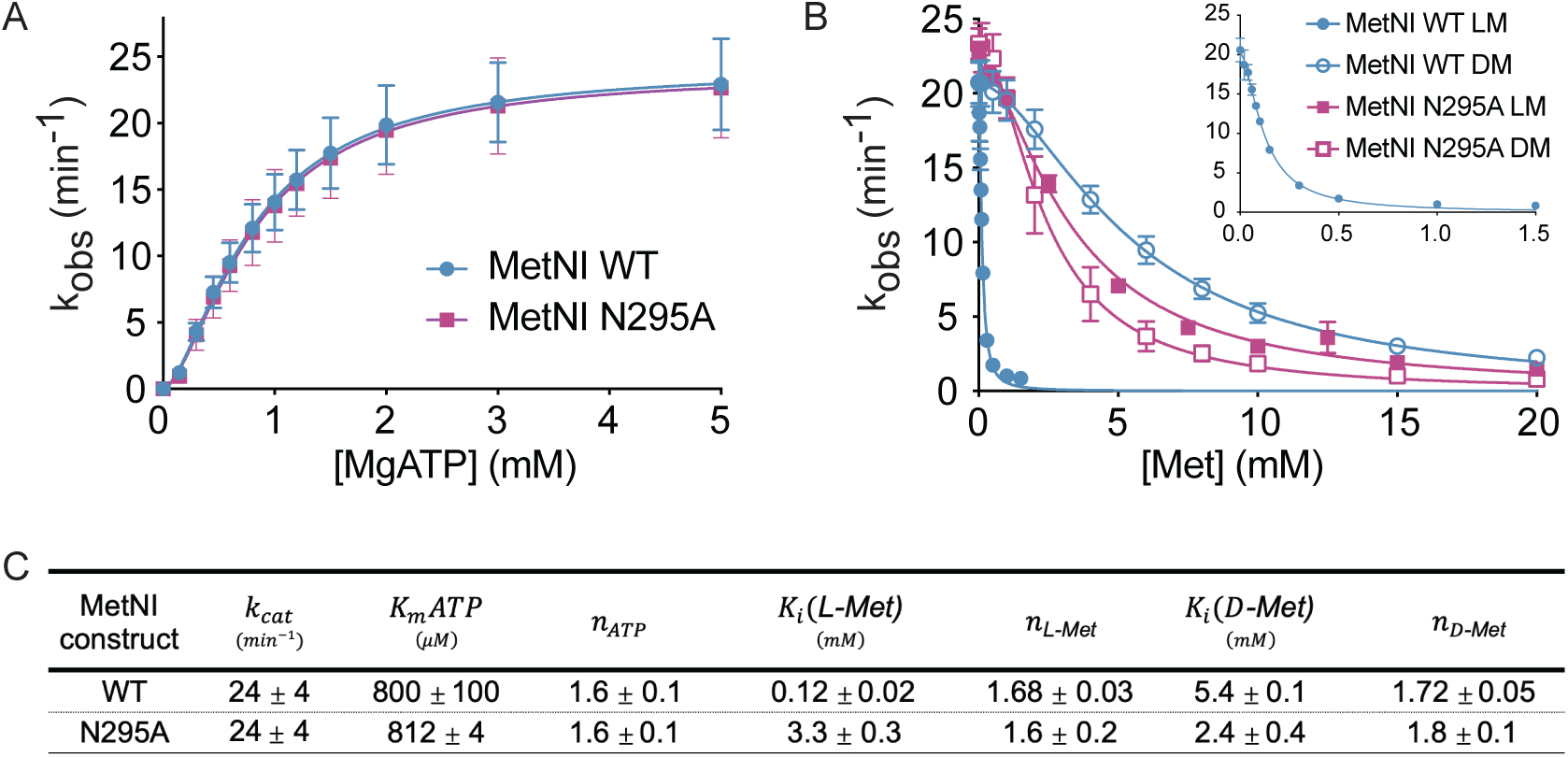
The MetNI N295A mutation neutralizes the trans-inhibitory effect of L-Met. (a) ATPase activity of WT MetNI (blue) and N295A MetNI (purple) as a function of MgATP. Reactions contained 500 nM MetNI solubilized in DDM. (b) Trans-inhibition of WT MetNI (blue) and N295A (purple) by L-Met (closed symbols) and D-Met (open symbols) in the presence of 5 mM MgATP. *Inset*: observed rate constants for WT MetNI at low concentrations of L-Met. (c) Summary of kinetic data from ATPase assays.

### Methionine binding by MetQ

To test if MetQ is capable of discriminating between methionine enantiomers, we measured the affinity between substrate and MetQ using isothermal titration calorimetry (ITC) (Fig. 3a). In all of the experiments described in this study, we utilized secreted (non-lipidated) MetQ (Fig. S2). Previous studies have shown that MetQ lipidation enhances its interaction with MetNI through association with the lipid bilayer (50, 51). For this work, we chose to use non-lipidated MetQ in order to reveal the underlying requirements for the intrinsic interaction with MetNI in the absence of co-localization to the membrane. Although MetQ lipidation will clearly influence the MetNI-Q system, meaningful interpretations can be made based on the relative activities within our non-lipidated system.

**Figure 3.**
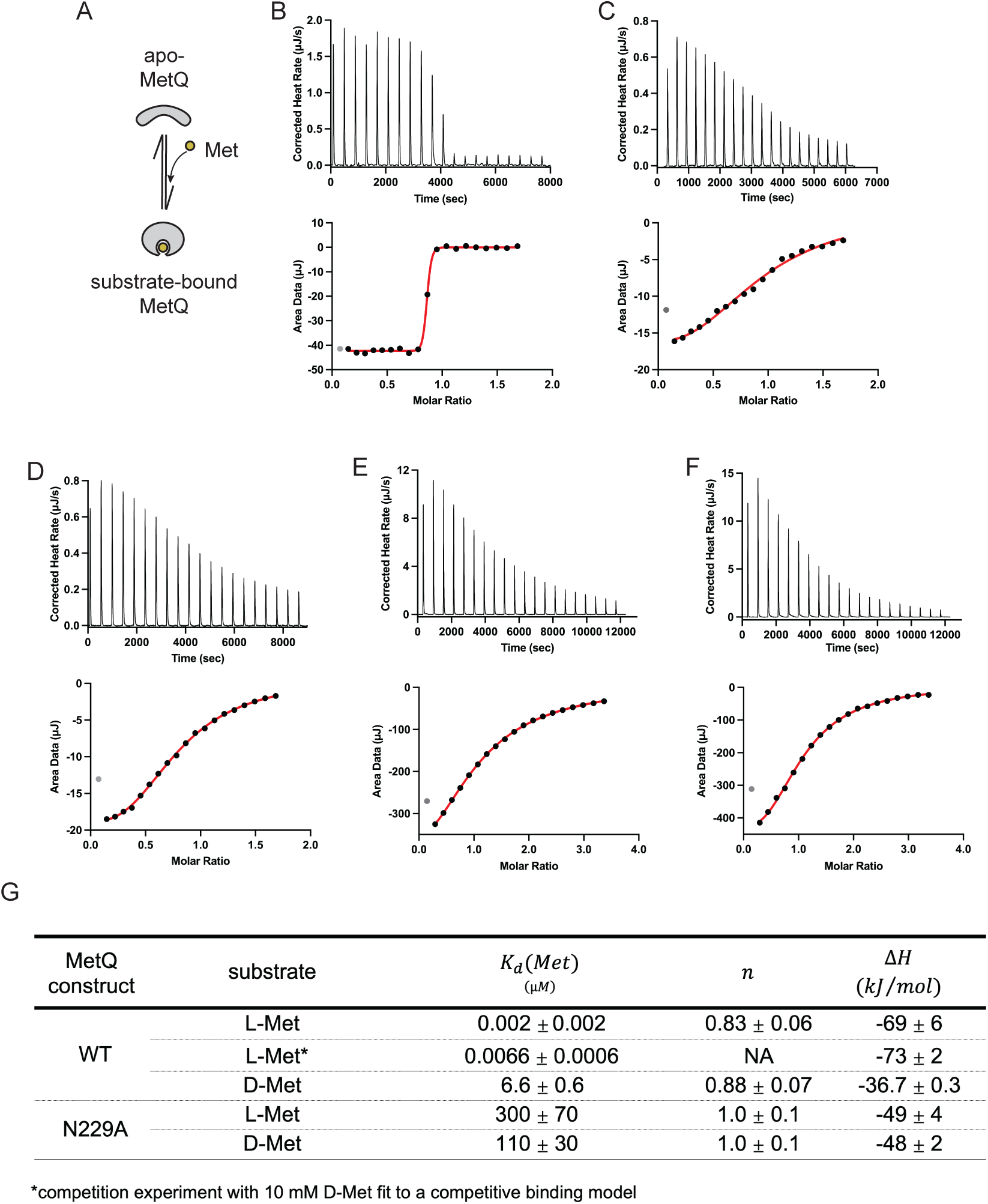
MetQ – substrate binding affinities. (a) Schematic of substrate binding by MetQ. (b-f) Representative isothermal titration calorimetry binding curves for (b) L-Met binding to WT MetQ, (c) L-Met competitive binding to WT MetQ bound to D-Met, (d) D-Met binding to WT MetQ, (e) L-Met binding to N229A MetQ, and (f) D-Met binding to N229A MetQ. Concentrations of protein and Met are given in the Materials and Methods. Gray datum point in each subpanel is excluded from the fit. (g) Dissociation constants derived from ITC titrations.

To study the affinity between methionine and wild-type MetQ, we first had to remove co-purifying L-Met via an established unfolding-refolding protocol (52, 53). Using this purified apo-MetQ, ITC experiments showed that wild-type MetQ binds L-Met with low nanomolar affinity (K_d_ = 6.6 ± 0.6 nM) (Figs. 3b-c). Furthermore, wild-type MetQ had a 1,000-fold stronger preference for L-Met over D-Met (K_d_ = 6.6 ± 0.6 µM) (Fig. 3d). To provide a form of MetQ that would serve as a constitutive apo-state mimic, we next tested the affinity of methionine for a previously-identified MetQ binding site mutant. This N229A mutation has been shown to both reduce MetQ affinity for D-Met and also facilitate D-SeMet transport *in vivo* (43). We determined that the N229A mutation reduced the affinity for L-Met by over five orders of magnitude (K_d_ = 300 ± 70 µM), similar to that for D-Met (K_d_ = 110 ± 30 µM) (Figs. 3e-f).

Based on these measurements, we demonstrated that MetQ can distinguish between L- and D-Met and very strongly selects for L-Met. We also concluded that the N229A mutant would be mostly unbound under the concentrations of L- and D-Met used in our experiments and could thus serve as a valid apo-MetQ mimic (Fig. 3g). Importantly, these K_d_ values allowed us to design subsequent experiments with WT MetQ in defined substrate-bound states – either apo, L-Met saturated, or D-Met saturated. Additionally, when possible, subsequent experiments were performed under identical buffer conditions as used in ITC experiments.

### Requirements for MetNI - MetQ complex formation

We next sought to investigate the substrate and nucleotide requirements for binding between MetNI and MetQ. To detect complex formation, we conducted fluorescence anisotropy experiments using fluorescein-labeled MetQ (Fig. 4a). The labeled MetQ variants were as follows: (i) wild-type native MetQ, containing copurified L-Met, denoted WT(LM), (ii) unfolded-refolded MetQ, denoted WT(apo), and (iii) unfolded-refolded MetQ with 100 µM D-Met, denoted WT(DM). For all experiments, we measured MetQ binding to a DDM-solubilized double mutant of MetNI. To capture the transporter in distinct nucleotide states, this double mutant featured an ATP hydrolysis-deficient E166Q Walker B motif mutation combined with the N295A inhibition mutation characterized above. For ease of reading, all references to MetNI in the anisotropy experiments described below indicate the use of the E166Q/N295A double mutant.

**Figure 4.**
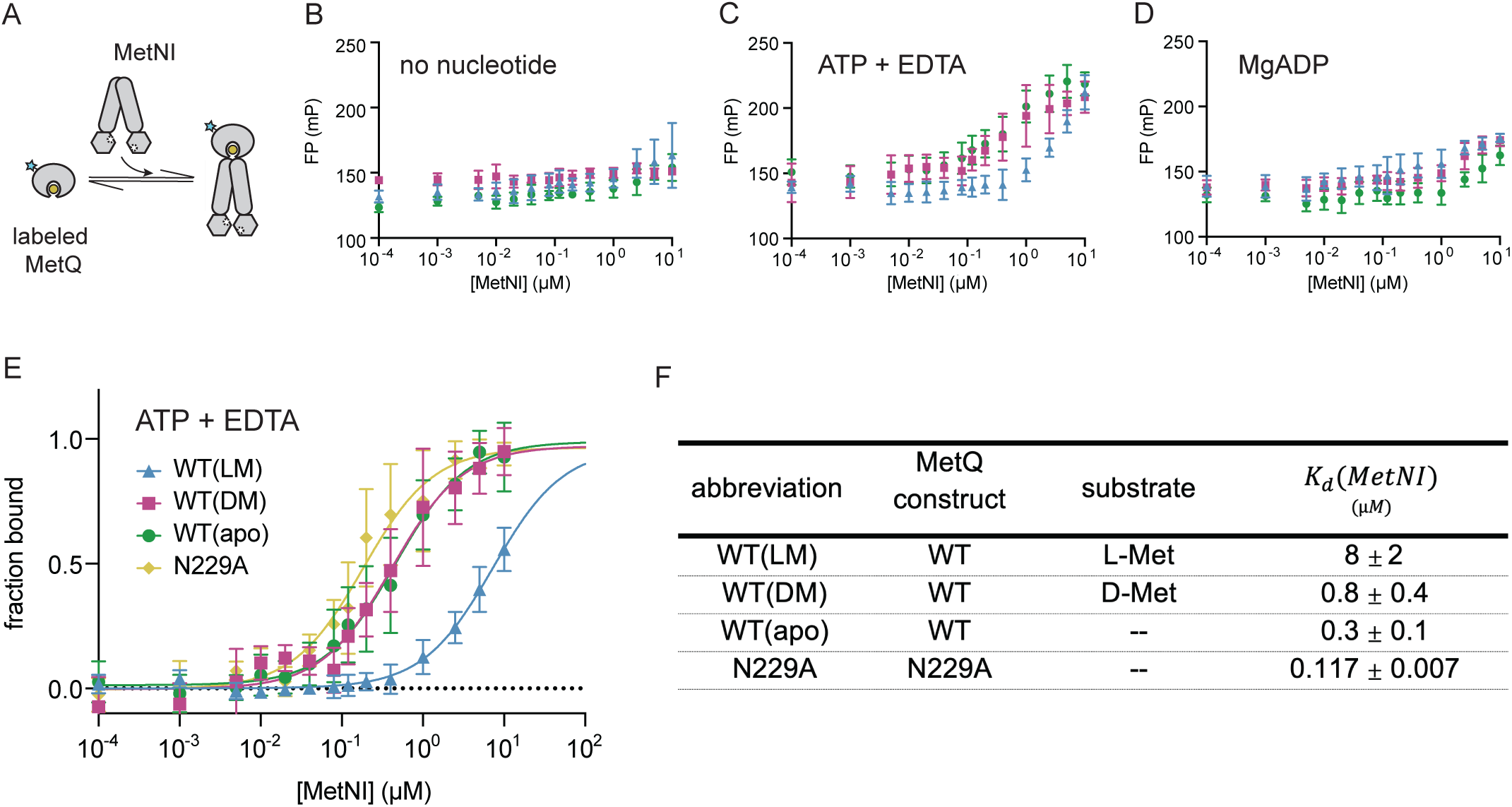
Effect of nucleotide-state and substrate on MetNI-Q complex formation. (a) Schematic of fluorescence anisotropy experiment. (b-d) Raw data for MetQ in (b) the absence of nucleotide, (c) presence of ATP/EDTA, and (d) MgADP. WT MetQ saturated with L-Met is shown in blue, saturated with D-Met in purple, and without substrate is in green. (e) Normalized binding curves for MetQ variants in the presence of ATP/EDTA. (f) Dissociation constants derived from anisotropy assays.

First, we examined binding between MetQ variants and MetNI as a function of nucleotide state. MetQ and MetNI did not form an observable complex in the absence of nucleotide or in the presence of MgADP, regardless of the presence or absence of methionine (Figs. 4a and 4c). In contrast, all MetQ variants formed a complex in the ATP-bound state, which was mimicked by the addition of ATP and EDTA (Fig. 4b). To compare the affinity of MetQ variants for MetNI in the ATP-bound state, we fit our data to a single-site binding model (see Materials and Methods, Figs. 4d and S3a). The substrate-free MetQ variants, WT(apo) and N229A, had stronger affinities for MetNI than WT(LM) (K_d_ = 0.6 ± 0.5 µM, 0.3 ± 0.2 µM, and 8 ± 2 µM, respectively, see Fig. 4e). Additionally, WT(DM) had a ∼10-fold tighter affinity for MetNI than WT(LM) (K_d_ = 0.5 ± 0.3 µM and 8 ± 2 µM, respectively). To verify that the unfolding-refolding process did not negatively affect the function of MetQ, we compared the K_d_ of native N229A and unfolded-refolded N229A and found them to be within error of each other (Fig. S3b).

Overall, our thermodynamic studies provide an initial equilibrium framework for methionine selectivity. ITC measurements show that MetQ preferentially binds L-Met over D-Met by three orders of magnitude. Our anisotropy experiments establish that binding of MetQ to MetNI is only detectable in the ATP-bound state. Furthermore, the affinity between MetQ and MetNI is the weakest in the presence of L-Met.

### MetQ stimulation of MetNI ATPase activity

With the thermodynamic parameters of the MetNI-Q system in hand, we next studied the kinetic properties of MetNI with MetQ variants (Fig. 5a). Studies on other ABC transporter systems show that detergent-solubilized ABC transporters exhibit significant uncoupled basal ATPase activity that can be stimulated by substrate and the SBP (54–56). To test if MetQ similarly stimulates MetNI, we first measured ATPase activity in detergent micelles (Fig. 5b). Under saturating MgATP concentrations, WT(apo) stimulated MetNI ATPase activity ∼10-fold above the basal rate (k_cat_ = 240 ± 30 min^-1^ and k_cat_ = 24 ± 2 min^-1^, respectively) while WT(LM) exhibited a marginal twofold stimulation effect (k_cat_ = 50 ± 20 min^-1^). WT(DM) also exhibited less pronounced stimulation, although at an intermediate level (k_cat_ = 110 ± 20 min^-1^). For the latter experiment, 100 µM exogenous D-Met was added to experiments to saturate MetQ with substrate. As a control to ensure that the exogenous Met did not interfere with the rate of hydrolysis, we compared the ATPase activity stimulated by native WT(LM) to that of unfolded-refolded WT MetQ with 100 µM exogenous L-Met. The resulting kinetic parameters were within error of each other (Fig. S4).

**Figure 5.**
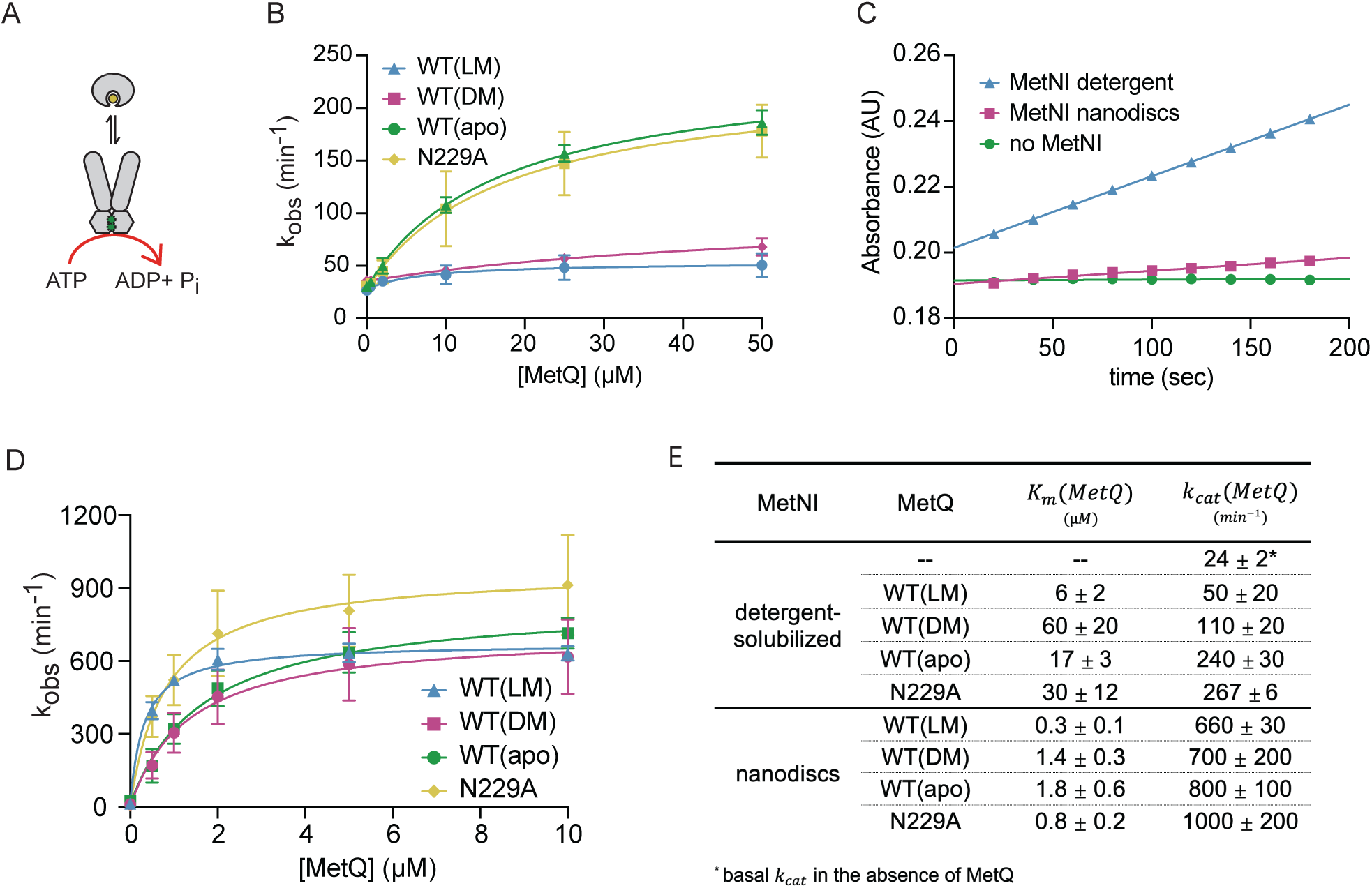
MetQ stimulates the ATPase activity of MetNI. (a) Schematic of stimulated ATP hydrolysis. (b) MetQ stimulation of detergent-solubilized N295A MetNI (100 nM) in the presence of 5 mM MgATP. (c) 200 nM N295A MetNI in DDM (blue) or reconstituted in nanodiscs (purple) was mixed with 5 mM MgATP in the absence of MetQ. A reaction without MetNI is shown in green. (d) MetQ stimulation of N295A MetNI nanodiscs (32 nM) in the presence of 5 mM MgATP. (e) Summary of kinetic data from ATPase assays.

The nature of the basal ATPase activity of ABC transporters is puzzling, as uncoupled activity is commonly perceived as a wasteful use of cellular energy. Similar observations in other systems suggest that futile cycling may be due to the non-native environment of detergent micelles (54, 57–59). To address this concern, we next reconstituted N295A MetNI into lipid bilayer nanodiscs. In comparison to detergent-solubilized MetNI, the nanodiscs showed trace levels of uncoupled activity over background (Fig. 5c). This suggests that the lipid bilayer may provide a more appropriate context for functional studies. Next, we added varying concentrations of MetQ to nanodiscs under saturating MgATP concentrations (Fig. 5d). Unlike that seen in detergent experiments, all MetQ variants robustly stimulated the ATPase activity of MetNI nanodiscs, including WT(LM) (Fig. 5e). MetQ-stimulated MetNI nanodiscs hydrolyzed ATP at levels approximately 5-10 times faster than that of detergent-solubilized MetNI. Additionally, the kinetic parameters measured in nanodiscs were only modestly different between MetQ variants; the K_m_ values ranged from 0.3 mM to 1.8 mM, and the k_cat_ values were 1.5-fold different. In summary, these kinetic data demonstrate the ability of MetQ to stimulate MetNI ATPase activity to comparable levels regardless of the presence of substrate.

### MetNI-Q activity in proteoliposomes

Finally, to monitor the translocation of methionine enantiomers across the lipid bilayer, we reconstituted MetNI into proteoliposomes. For all liposomal studies, we utilized wild-type MetNI in order to more accurately replicate *in vivo* conditions. Furthermore, we did not make any assumptions regarding the orientation of MetNI when inserted into the membrane (Fig. 6a). We first measured the ATPase activity of MetNI in proteoliposomes in the presence of different MetQ variants. For these experiments, saturating concentrations of methionine-bound MetQ (50 µM) were incorporated into the lumen of proteoliposomes. The reaction was initiated by adding saturating MgATP to the outside of the proteoliposomes (Fig. 6a). As seen in nanodiscs, there was very little uncoupled ATPase activity in proteoliposomes. Interestingly, WT(apo), WT(LM), and WT(DM) stimulated MetNI equally, and the specific activities were within error of each other (23 ± 4, 23 ± 2, and 18 ± 7 nmol P_i_ · min^-1^ · mg^-1^ MetNI, respectively, see Fig. 6b). We also tested the MetQ stimulated ATPase activity using N229A MetQ (Fig. 6c). The rates of ATP hydrolysis were very similar in the absence and present of substrate; however, unlike in nanodiscs, N229A MetQ stimulated rates in proteoliposomes were half of the WT MetQ stimulated rates (Fig. 6g).

**Figure 6.**
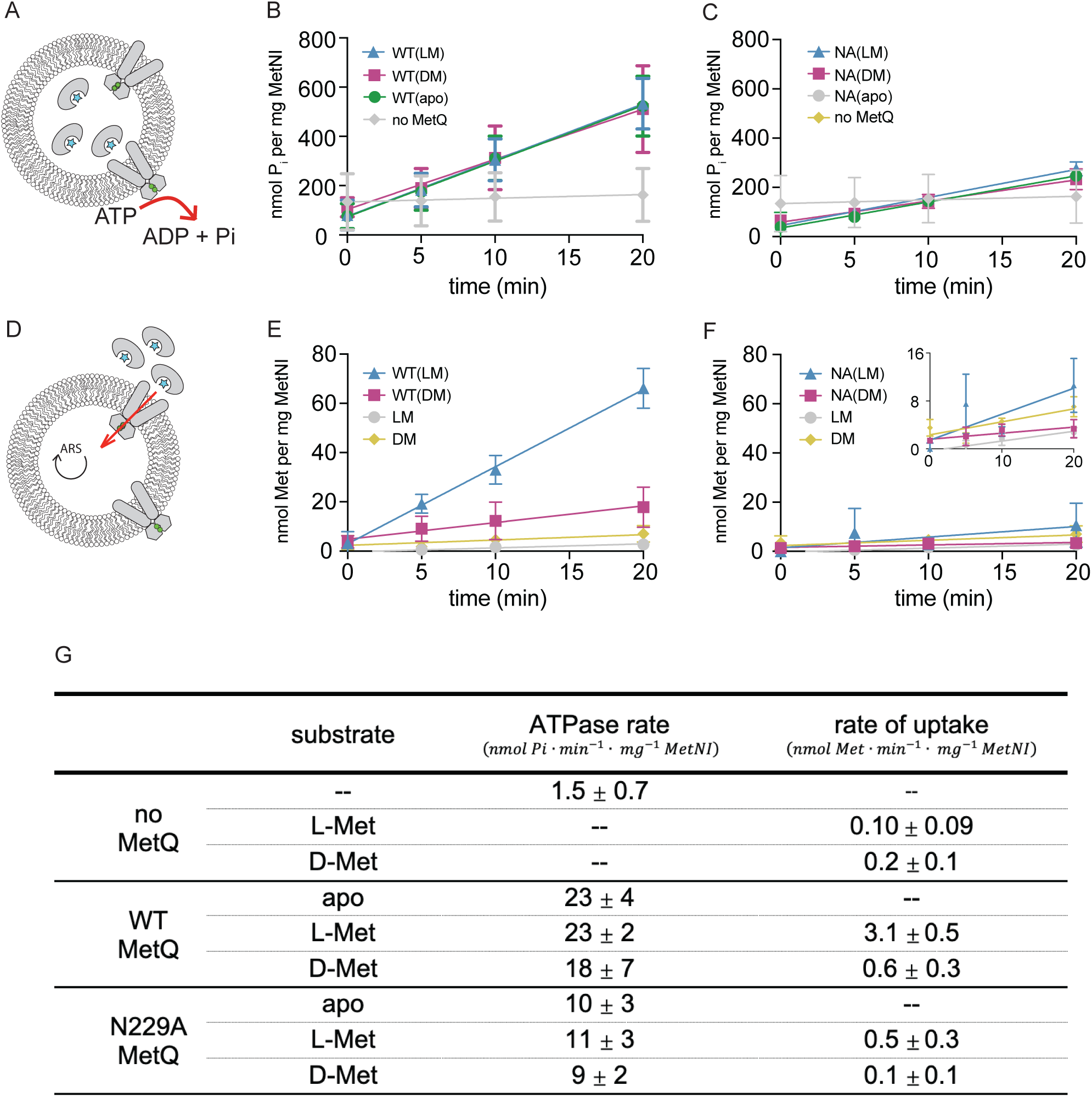
Transport and ATPase activity of wild-type MetNI proteoliposomes. (a) Schematic of ATPase assay. (b and c) ATP hydrolysis in the presence of WT MetQ (b) and N229A MetQ (c) with and without substrate. (d) Schematic of transport assay. (e and f) Substrate transport in the presence of WT MetQ (e) and N229A MetQ (f). *Inset* in (f) shows data at low specific activity values. (g) Summary of kinetic data from proteoliposomes.

Lastly, to detect the transport of substrate into proteoliposomes, we first incorporated an ATP-regeneration system in the lumen (Fig. 6d). Next, we prepared a stock of each enantiomer combined with a trace of amount of the identical ^3^H enantiomer. MetQ was then incubated with the substrate, and the reaction was initiated by adding the MetQ-methionine mixture to the proteoliposomes. Under saturating conditions, L-Met was imported into proteoliposomes with an uptake rate of 3.1 ± 0.5 nmol L-Met · min^-1^ · mg^-1^ MetNI (Fig. 6e). Based on estimates of the internal volume of 400 nm proteoliposomes, L-Met was transported against the concentration gradient (60). Despite saturating concentrations and similar ATPase rates, D-Met was transported 5-fold more slowly (0.6 ± 0.3 nmol D-Met · min^-1^ · mg^-1^ MetNI). Lastly, we tested the ability of N229A MetQ to support methionine transport (Fig. 6f). We found that L-Met was transported five times slower with N229A than with WT MetQ, and uptake of D-Met above background was not detected.

From these transport assays we conclude that the MetNI transporter, in addition to the SBP, plays a role in substrate selectivity. The molecular basis of this discrimination does not simply rely on the stimulation of ATP hydrolysis, as the ATPase rate is equal in the absence or presence of L- or D-Met.

## Discussion

In this work, we quantitatively examine individual steps of the methionine import cycle. Our work suggests that the mechanistic basis of substrate preference relies on both the MetQ SBP and the MetNI transporter, which work in concert to select for import of L-Met over D-Met. Our model for methionine selectivity is illustrated in Figure 7 and discussed below.

**Figure 7.**
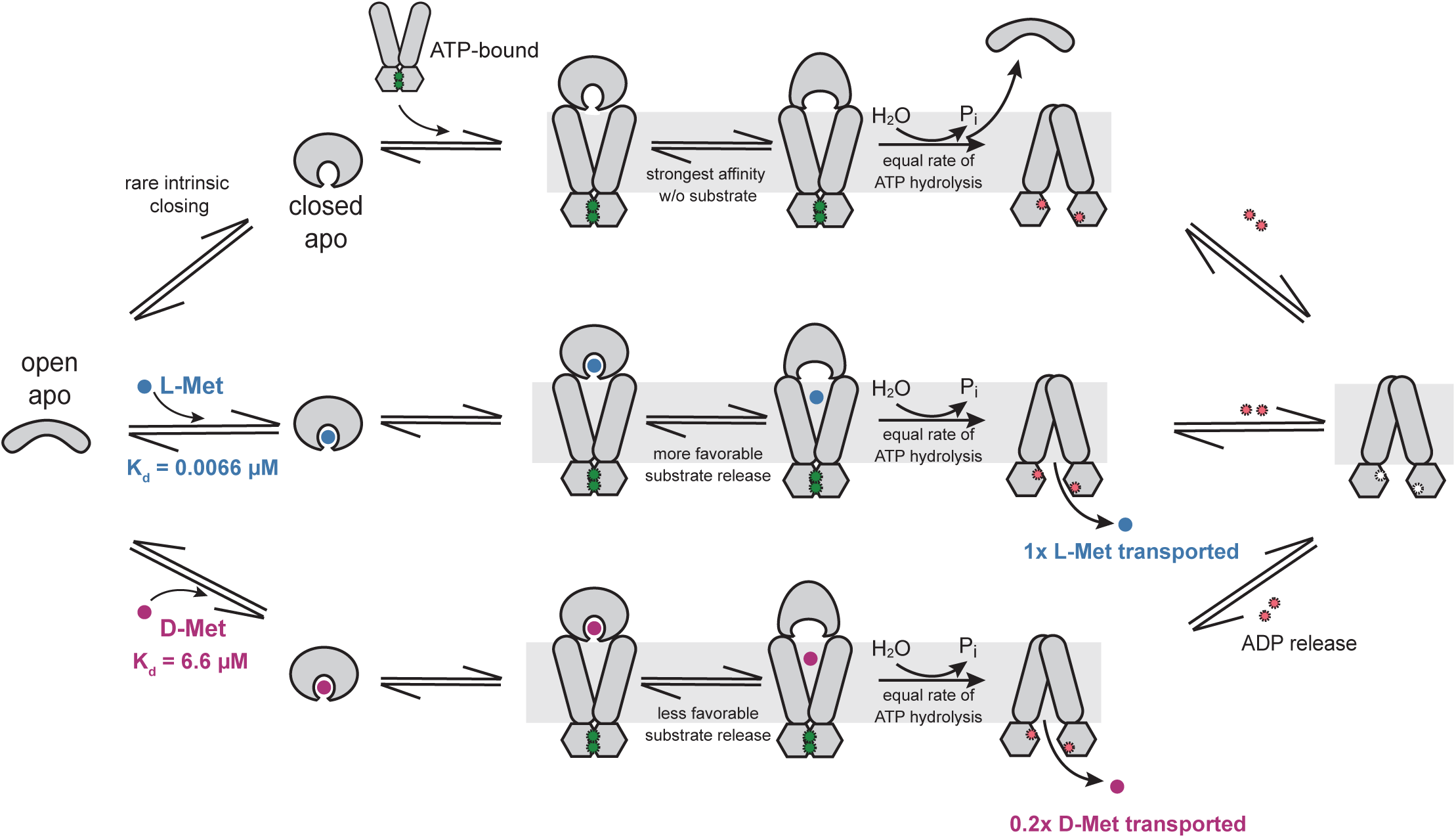
Discrimination between L- and D-Met by the MetNI-Q transport system. This model incorporates insights from this work as well as multiple single molecule, structural, and functional studies of related ABC importers. (Top pathway) MetQ in the open-apo state infrequently samples the closed-apo conformation, referred to as MetQ(apo). This rarely-occurring MetQ(apo) species can form a complex with outward-facing ATP-bound MetNI. Complex formation drives a conformational change in MetQ to a “twisted” state. MetNI then hydrolyzes ATP and adopts the inward-facing conformation, leading to MetQ and P_i_ dissociation. ADP is subsequently released from MetNI. (Middle pathway) MetQ in the open-apo state binds L-Met with very tight affinity, inducing the closed substrate-bound state MetQ(LM). Similar to the top pathway, MetQ(LM) forms a complex with ATP-bound MetNI. In a concurrent step, MetQ(LM) adopts the twisted state and releases the substrate to the transmembrane cavity. The release of substrate is energetically favorable, given that the affinity of ATP-bound MetNI is stronger for MetQ(apo) than for MetQ(LM). ATP hydrolysis triggers rearrangement to the inward-facing conformation. L-Met is transported across the membrane and MetQ, P_i_, and ADP are released from MetNI before another round of transport can commence. (Bottom pathway) MetQ in the open-apo state binds D-Met with 1,000-fold weaker affinity than L-Met. Upon MetQ(DM) docking to MetNI, D-Met is released from MetQ; however, the release is less favorable than with L-Met given the intermediate affinity between ATP-bound MetNI and MetQ(DM). Subsequent steps are the same as for L-Met transport.

Our ITC measurements reveal that MetQ plays a significant role in selectivity and has a 1,000-fold preference for L-Met over D-Met. The nature of this disparate recognition is currently unclear. Despite vastly different binding affinities, the crystal structures of *N. meningitides* MetQ bound to methionine depict nearly identical side chain interactions with the L- and D-enantiomers (52). In the future, it will be of particular interest to study the dynamics of MetQ in solution with different substrates, as done with other SBPs using smFRET and EPR (30–33). Furthermore, structural studies indicate that MetQ employs the canonical hinge-type interconversion between open-apo and closed-bound conformations. However, unlike other transporters, MetQ adopts a third distinct conformation; it forms a unique “twist” conformation when in complex with the transporter (47). How the identity of the substrate affects adoption of the twist conformation and how this in turn affects the rate of substrate dissociation will be of great interest to investigate in the future.

Our evaluation of MetNI-Q parameters in different lipid environments reveal that while the absolute values of the kinetic constants are unequal, trends between the MetQ variants are consistent. For example, in both detergent and nanodiscs the K_m_(MetQ) value for WT(LM) is lower than the other MetQ variants. Furthermore, the k_cat_ values for ATPase stimulation in both detergent and nanodiscs are ordered WT(apo) > WT(DM) > WT(LM), albeit the differences in nanodiscs are very narrow (Fig. 5d). In line with the latter, we find that the k_cat_ for MetQ variants in proteoliposomes are within error of each other (Fig. 6g). Overall, these experiments highlight the importance of considering the lipid environment when interpreting any ABC transporter data.

The *in vitro* transport rates for L- and D-Met measured here (3.1 and 0.6 nmol/min/mg protein, respectively) are comparable to that found in other reconstituted proteoliposome systems (0.11 – 1.2 nmol/min/mg protein) and are within reason of those measured *in vivo* for D-SeMet (6.3 nmol/min/mg protein) (47, 54, 61, 62). Our specific activity measurements do not take into account the vectorial reconstitution ratio of the transporter, and thus it is not possible to calculate the turnover number or stoichiometry of ATP molecules hydrolyzed per methionine transported. Further studies will require a more complete investigation of the kinetic and thermodynamic parameters in proteoliposomes. Lastly, we note that N229A MetQ supported transport is low relative to WT MetQ, unlike that seen in published studies (47).

The two most unanticipated results from this study are that: (i) apo-MetQ fully stimulates ATPase activity and (ii) D-Met transport is slower than L-Met, even in the presence of saturating MetQ and methionine concentrations. With regard to (i), this finding upends the assumption that substrate is required for maximal transporter ATPase activity. Stimulation by an apo-SBP has been observed in other liposomal systems; however, in these cases, the apo-SBP stimulates the transporter noticeably less than the substrate-bound form (63, 64). Here we see that the docking of apo-MetQ robustly activates MetNI for transport, indicating that a closed conformation of apo-MetQ is sufficient for full stimulation. Based on previous work on SBPs, we infer that the occurrence of the closed apo-MetQ conformation is very rare, and how this state may influence overall transport rates is not yet apparent.

Regarding (ii), the different rates of L-Met and D-Met transport can be rationalized by considering the relative affinities of WT(LM), WT(DM), and WT(apo) for ATP-bound MetNI. WT(apo) has a ∼25-fold stronger interaction with MetNI than WT(LM) (K_d_ = 0.3 µM and 8 µM, respectively). The tighter affinity of WT(apo) for MetNI supports a model in which release of L-Met from the MetNI-WT(LM) complex is energetically favorable. In this model, the energetic “cost” of substrate release from MetNI-WT(LM) is compensated for by the increased binding interactions between the newly-formed WT(apo) and MetNI. In turn, L-Met dissociation from MetQ into the TMD permeation pathway allows for a productive transport cycle. The reduced rate of D-Met transport relative to L-Met is consistent with this model; the more subtle difference between the dissociation constants of WT(apo) and WT(DM) for ATP-bound MetNI suggests that the release of D-Met from MetQ is less favored in relation to L-Met (K_d_ = 0.3 µM and 0.8 µM, respectively). We propose that this difference in methionine release accounts in part for the observed difference in L-Met and D-Met transport rates. More favorable L-Met release yields a higher transport rate, while less favorable D-Met release correlates with slower transport. In summary, our findings demonstrate how MetNI-Q complex formation provides an additional stage for substrate discrimination.

Together, these quantitative analyses lay a foundation for understanding the molecular basis of substrate selectivity in ABC transporters. The MetNI-Q system exhibits multiple distinct mechanisms for discrimination that together dictate the preference for L-Met over D-Met. The incorporation of multiple mechanisms raises the possibility that ABC transporters may each utilize a different combination of mechanisms to fine tune substrate specificity. This versatility would allow for both optimal growth in abundant conditions as well as survival in limiting environments.

## Experimental procedures

### Protein expression and purification

MetNI was prepared as previously described (48). Expression and purification of MetQ was carried out as described with minor modifications (45). Briefly, MetQ with a C-terminal 6x-histidine tag was expressed in *E. coli* BL21 (DE3) Gold cells (Agilent) grown at 37 °C in Terrific Broth with 100 µg/ml ampicillin. Cells were induced at OD_600_ 2.0 with 1 mM IPTG for one hour, harvested via centrifugation, and stored at −80 °C.

To isolate the periplasmic extract containing secreted non-lipidated MetQ, 10 g of cells were resuspended in 10 mL of 10 mM Tris pH 7.5, 150 mM NaCl, and 40% sucrose and stirred for 1 hour at room temperature. Cells were then osmotically shocked by the addition of 500 mL of ice-cold water and stirred for 10 minutes at 4 °C. Following osmotic shock, buffer components were added to the periplasmic extract to a final concentration of 25 mM Tris pH 7.5, 150 mM NaCl, and 5 mM β-mercaptoethanol (*β*ME). After centrifugation at 44,000 x g for 30 minutes at 4 °C, 200 µg/ml DNase I was added to the supernatant and incubated on ice for 20 minutes. The supernatant was then filtered through a 0.65 µm filter and imidazole added to a final concentration of 17 mM. The periplasmic extract was injected onto a 5 mL HisTrap HP column (Cytiva) and washed with MetQ buffer (25 mM Tris pH 7.5, 150 mM NaCl, 5 mM *β*ME) supplemented with 17 mM imidazole for 5 column volumes. MetQ was eluted using MetQ buffer containing 400 mM imidazole and buffer exchanged into MetQ buffer using a HiPrep 26/10 desalting column (Cytiva). The eluent was then injected on a HiLoad 16/600 Superdex 200 pg column equilibrated in MetQ buffer (Cytiva). Peak fractions were concentrated to ∼ 1.5 mM using a 10 kD MWCO filter (Millipore), and aliquots were flash frozen in liquid nitrogen and stored at −80 °C.

To purify substrate-free MetQ, protein bound to the HisTrap column was unfolded using 10 column volumes of unfolding buffer (25 mM Tris-HCl pH 7.5, 150 mM NaCl, 400 mM, 5 mM *β*ME, and 6 M guanidine-HCl) at a flow rate of 1.5 mL/min. To refold MetQ, a linear gradient from 100% to 0% unfolding buffer was flowed at 1.0 mL/min over 12 column volumes, followed by an additional 10 column volumes of MetQ buffer to ensure complete refolding. Elution from the HisTrap column and subsequent purification steps were performed as described above.

### MetQ labeling

MetQ was labeled at the single native cysteine at amino acid position 23. One mL of ∼350 µM purified MetQ was buffer exchanged into labeling buffer (25 mM Tris pH 7.0, 150 mM NaCl, 5 mM EDTA, 10 mM TCEP) using a 5-mL HP Desalting column (Cytiva) at 4 °C. Peak fractions were pooled, and labeling buffer was added for a final MetQ concentration between 30-40 μM. Solid fluorescein 5-maleimide (Thermo Fisher) was added directly to MetQ to a final concentration of 1 mM. The solution was continuously inverted at room temperature for 2 hours. The labeling reaction was then quenched by adding *β*ME to a final concentration of 100 mM and inverted at room temperature for an additional 15 minutes. Next, the solution was flowed over a column containing 200 μL of Ni-NTA beads (Qiagen) equilibrated with MetQ buffer at room temperature. The column was washed with 10 mL of MetQ buffer before elution with MetQ buffer containing 400 mM imidazole. The eluted protein was exchanged into MetQ buffer using a 5 mL HP Desalting column at 4 °C. The peak fractions were pooled, and aliquots were flash frozen in liquid nitrogen and stored at −80 °C.

### Reconstitution of MetNI into lipid bilayers

MetNI nanodiscs were prepared as previously described (65). Briefly, egg L-phosphatidylcholine and *E. coli* polar lipid extract (Avanti Polar Lipids, Inc.) were mixed in a 1:3 (w/w) ratio, dried, and resuspended in lipid buffer (50 mM TAPS pH 8.5, and 250 mM NaCl). Purified MetNI, MSP1D1(-) (Sigma Aldrich), and resuspended lipids were combined at a 1:3:60 molar ratio, and DDM was added to 12 mM final. The mixture was rotated at 4 °C for one hour. Following removal of detergent using 0.6 g/ml Bio-Beads SM2 resin (Bio-Rad) for three hours, MetNI nanodiscs were purified using a 1-mL HisTrap HP column (Cytiva) followed by injection onto a Superdex 200 Increase 10/300 column (Cytiva). Both columns were equilibrated in lipid buffer. Peak fractions were pooled, and aliquots were flash frozen in liquid nitrogen and stored at −80 °C. The concentration and stoichiometry of MetNI nanodiscs was determined using gel densitometry.

MetNI proteoliposomes were prepared as previously described with minor modifications (60). Briefly, 1:3 (w/w) egg PC:*E.coli* polar lipid extract in lipid buffer was diluted to 4 mg/ml and destabilized with 0.3% Triton X-100 final for 20 minutes at room temperature. Purified MetNI was mixed with destabilized liposomes at a 1:50 (w/w) ratio and incubated for 20 minutes at room temperature. Following four rounds of detergent removal using Bio-Beads, MetNI liposomes were harvested via centrifugation at 267,000 x g for 20 minutes at 4 °C. The pellet was resuspended to 20 mg/ml lipids in liposome buffer (25 mM Tris pH 7.5 and 150 mM NaCl). Proteoliposomes were harvested and resuspended again in liposome buffer. Aliquots were flash frozen in liquid nitrogen and stored at −80 °C. The concentration of MetNI in liposomes was determined using gel densitometry and ranged from 0.3 – 0.7 µM (Fig. S5).

### Reporting of thermodynamic and kinetic parameters

For all assays, parameters were determined for each experimental trial independently, and the average values across the trials were reported with the standard deviation. Error bars represent standard deviation. At least three independent experiments were performed using at least two separate preparations of each protein.

### ATPase assays

For assays with detergent-solubilized MetNI, the amount of inorganic phosphate generated in solution was measured in real time using the EnzChek Phosphate Assay Kit (Thermo Fisher). For assays without MetQ, each 100 µl reaction contained 500 nM DDM-solubilized MetNI, 57.5 mM Tris pH 7.5, 10 mM TAPS pH 8.5, 150 mM NaCl, 0.05% DDM, 200 µM 2-amino-6-mercaptio-7-methylpurine riboside, 0.1 units of purine nucleoside phosphorylase, and L- or D-Met as indicated. Equimolar amounts of MgCl_2_ and ATP were manually added and mixed to initiate the reaction. Reaction components were incubated for 6 minutes at 37 °C before the start of the reaction, and the absorbance at 360 nm was measured every 20 seconds on an Infinite M200 microplate reader (Tecan Group) at 37 °C. Initial rates were obtained by calculating the linear portion of the change as a function of time. Data were fit as previously described (48). ATPase assays with varying MetQ were performed identically with 100 nM MetNI and 5 mM MgATP.

Nanodiscs ATPase assays were conducted using procedures similar to that described above. Each 100 µl reaction contained 32 nM MetNI nanodiscs, 57.5 mM Tris pH 7.5, 10 mM TAPS pH 8.5, 150 mM NaCl, 1.5 mM *β*ME, 200 µM 2-amino-6-mercaptio-7-methylpurine riboside, and 0.1 units of purine nucleoside phosphorylase. 5 mM MgCl_2_, 5 mM ATP, and varying amounts of MetQ were manually mixed with nanodiscs to initiate the reaction.

For both nanodiscs and detergent-based ATPase stimulation experiments, the observed rate constants were plotted as a function of MetQ concentration and fit to the following equation:

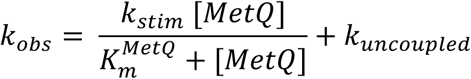

and

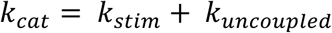

where *k*_*obs*_ is the observed rate constant, *k*_*stim*_ is the MetQ-stimulated rate constant, 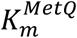 is the concentration of MetQ at which *k*_*obs*_ is equal to half of *k*_*stim*_, *k*_*uncoupled*_ is the observed rate constant in the absence of MetQ, and *k*_*cat*_ is the MetQ-stimulated maximal rate constant.

For ATPase assays using MetNI proteoliposome assays, MetQ, L-Met, and/or D-Met were incorporated into the lumen via three freeze-thaw cycles using liquid nitrogen. Following 11 rounds of extrusion through a 400 nm filter, proteoliposomes were harvested 267,000 x g for 20 minutes at 4 °C. Pellets were resuspended in liposome buffer and harvested again via centrifugation. Finally, proteoliposomes were resuspended to 5 mg/ml lipids in liposome buffer. Proteoliposomes and MgATP were separately incubated for 3 minutes at 37 °C and then mixed together to start the reaction. Final reaction conditions were 4 mg/ml lipids, 25 mM Tris pH 7.5, 150 mM NaCl, 10 mM MgCl_2_, 2 mM ATP, 0.06 – 0.14 µM MetNI, 50 µM L- or D-Met, and 50 µM MetQ. At given time points, 50 µl of the reaction was quenched with an equal volume of 12% SDS. The amount of inorganic phosphate generated was measured using a molybdate colorimetric method (66).

### Anisotropy studies

To determine the binding constant between MetNI and MetQ, the labeled MetQ concentration was held constant at 20 nM and the detergent-solubilized MetNI concentration varied from 0-10 μM. The final buffer conditions for anisotropy experiments were 57.5 mM Tris pH 7.5, 10 mM TAPS pH 8.5, 150 mM NaCl, 1.5 mM *β*ME, and 0.05% DDM. The ATP-bound state was mimicked with 5 mM ATP and 10 mM EDTA, and the measurements in the ADP-state included 5 mM MgADP.

For each trial, raw fluorescence polarization (FP) values and their corresponding MetNI concentrations were plotted and analyzed to determine the *K*_*d*_ using the equation:

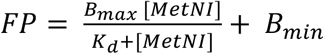

where *FP* is the raw fluorescence polarization value, *B*_*max*_ is the *FP* upper bound, *B*_*min*_ is the *FP* lower bound, and *K*_*d*_ is the MetNI concentration at which 50% of the maximum change in *FP* has been reached.

For each trial, raw FP values were converted to fraction bound values using the equation:

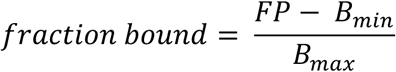

The average fraction bound value across trials was plotted as a function of MetNI concentration and fit to the following equation:

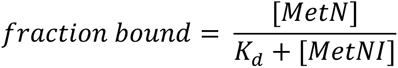

### Isothermal titration calorimetry

MetQ variants were dialyzed overnight into dialysis buffer (57.5 mM Tris pH 7.5, 10 mM TAPS pH 8.5, and 150 mM NaCl) using a Slide–A-Lyzer 10 kD MWCO MINI dialysis device (Thermo Scientific). The next day, samples were recovered and cleared at 313,000 x g for 20 minutes at 4 °C. Solid L- and D-Met (Sigma) were reconstituted in the dialysate. Binding data were collected on a Nano ITC (TA Instruments) at 25 °C. Each experiment consisted of 20 injections of 2.5 µL each, and the sample cell was allowed to equilibrate for 300 - 600 seconds between injections. 50 µM wild-type MetQ was titrated with 250 µM L- or D-Met, and 500 µM N229A MetQ was titrated with 5 mM L- or D-Met. For the competitive assay, MetQ was dialyzed overnight into dialysis buffer supplemented with 10 mM D-Met. The next day, samples were recovered and the concentration adjusted to 50 µM MetQ using the dialysate. L-Met was prepared in the dialysate to a final concentration of 250 µM, and injections were performed as described above. Each experimental trial was processed using the Independent or Competitive Replacement model on NanoAnalyze software.

### Transport assay

Transport assays were conducted as described with modifications (67). An ATP-regenerating system (ARS) was incorporated into the lumen of proteoliposomes for transport assays. 3x ARS (20 mM Tris pH 7.5, 150 mM NaCl, 15 mM MgCl_2_, 6 mM ATP, 72 mM creatine phosphate, and 7.2 mg/ml creatine kinase) was mixed in a 1:2 (v/v) ratio with proteoliposomes via three freeze-thaw cycles using liquid nitrogen. Proteoliposomes were then extruded 11 times through a 400 nm filter and harvested at 267,000 x g for 20 minutes at 4 °C. Pellets were resuspended in 20 mM Tris pH 7.5, 200 mM NaCl, and 5 mM MgCl_2_, harvested via centrifugation and resuspended again to 5 mg/ml final. A 1:50 mixture of [^3^H] L-Met (American Radiolabeled Chemicals, Inc.) to unlabeled L-Met was prepared in 20 mM Tris pH 7.5, 200 mM NaCl, and 5 mM MgCl_2_ and mixed with MetQ. The same mixture was prepared for D-Met experiments with the appropriate change in labeled and unlabeled enantiomer.

Proteoliposomes and Met-loaded MetQ were separately incubated for 3 minutes at 37 °C and then mixed together to start the reaction. Final reaction conditions were 4 mg/ml lipids, 20 mM Tris pH 7.5, 200 mM NaCl, 5 mM MgCl_2_, 0.06 – 0.26 µM MetNI, 50 µM L- or D-Met, and 50 µM MetQ. At indicated time points, 50 µl of the reaction was mixed with 300 µl of cold stop buffer (20 mM Tris pH 7.5, 200 mM NaCl, 5 mM MgCl_2_, 8% PEG 6000, and 200 µM unlabeled L- or D-Met). Quenched time points were filtered through a 96-well vacuum manifold filtration plate (Millipore) pre-equilibrated in stop buffer and washed twice with 200 µl stop buffer. After drying overnight, 250 µl scintillation fluid was added to each well, incubated for 15 minutes, and counted using a Perkin Elmer 1450 Wallac MicroBeta Trilux counter.

## Supporting information

This article contains supporting information.

## Acknowledgements

We gratefully acknowledge Geeta Narlikar for discussion and critical reading of the manuscript, Douglas Rees for the kind gifts of plasmids, the Narlikar group for assistance with anisotropy experiments, Michael J. Stevenson and the Stevenson group for assistance with ITC experiments, and Jeff Oda and Matt Helm for instrument support.

## Funding and additional information

This work was supported by the National Institutes of Health 5SC3GM144189 (to J.G.Y.). The content is solely the responsibility of the authors and does not necessarily represent the official views of the National Institutes of Health.

## Abbreviations

ABC: ATP-binding cassette
NBDs: nucleotide-binding domain
TMDs: transmembrane domains
SBP: substrate binding protein
SBD: substrate binding domain
ITC: isothermal titration calorimetry
DDM: n-dodecyl-*β*-D-maltoside
FP: fluorescence polarization
ARS: ATP-regenerating system
*β*ME: beta-mercaptoethanol.

