## Supporting information for "Two stages of substrate discrimination dictate selectivity in the *E. coli* MetNI-Q ABC transporter system"

Supporting figures 1-5

### Figure S1

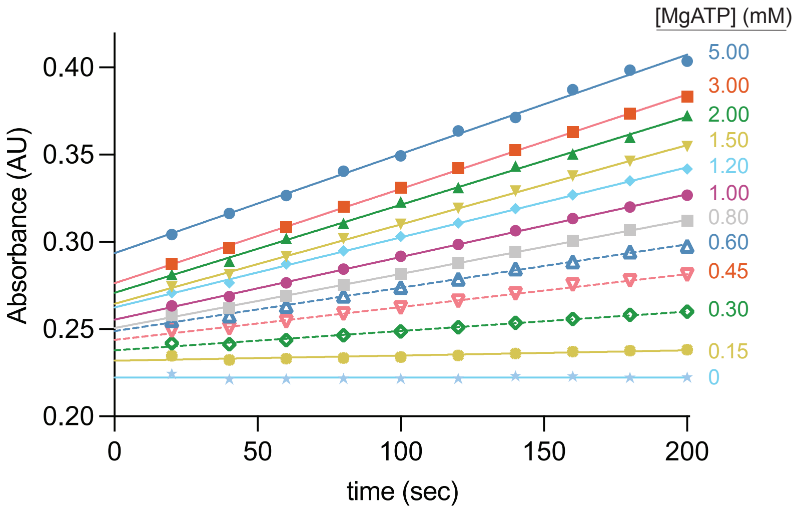


**Supporting Figure 1. Representative raw data from ATPase assays.** MgATP was mixed with detergent-solubilized MetNI and the absorbance at 360 nm was measured for 200 seconds. Reactions contained a final concentration of 500 nM MetNI and the indicated final concentration of MgATP.

#### Figure S2

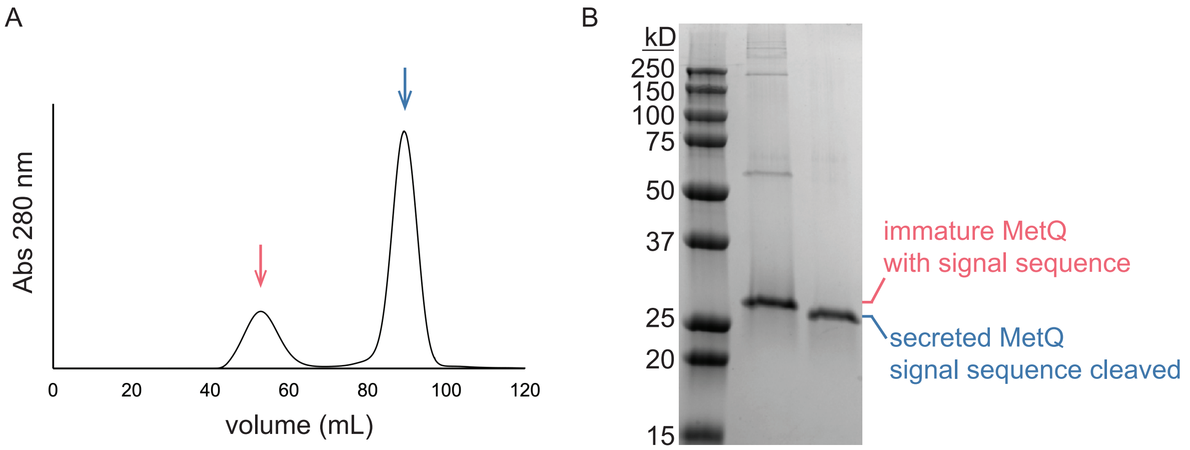


**Supporting Figure 2. Purification of secreted MetQ.** (a) Chromatogram of Ni-affinity purified His-MetQ injected onto a HiLoad 16/600 Superdex 200 pg column. The red arrow indicates insoluble material eluting at the void volume. The blue arrow highlights a monodisperse peak at ~90 ml elution volume. (b) SDS-PAGE of SEC fractions. Secreted MetQ was used in all experiments in this study.

#### Figure S3

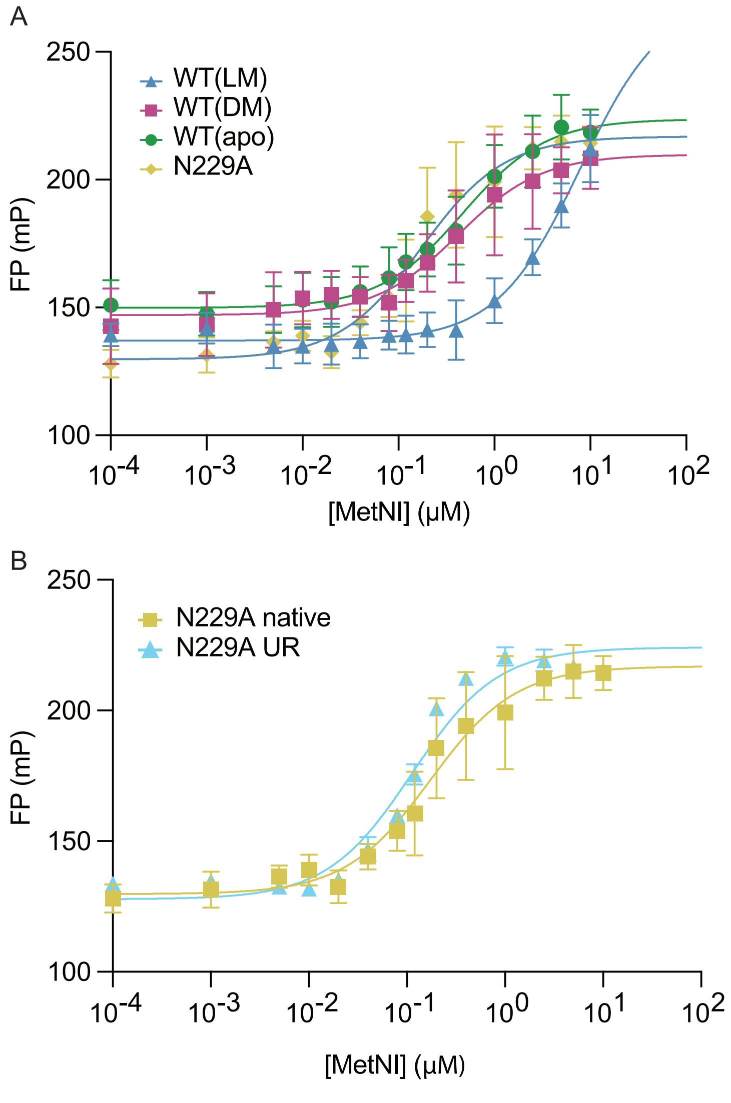


**Supporting Figure 3. Raw data from anisotropy experiments.** MetNI-Q complex formation as a function of substrate. Reactions contain 20 nM labeled MetQ**,** DDM-solubilized E166Q/N295A MetNI**,** and ATP/EDTA. (a) Raw data corresponding to Figure 4d in the main text. (b) Comparison of purified native N229A (yellow) and unfolded-refolded (UR) N229A (cyan). $K_d$ values are 0.3 ± 0.2 μM and 0.114 ± 0.008 μM, respectively. Data were fit to a one-site specific binding model to determine the $K_d$ and maximum and minimum FP values (see Materials and Methods).

#### Figure S4

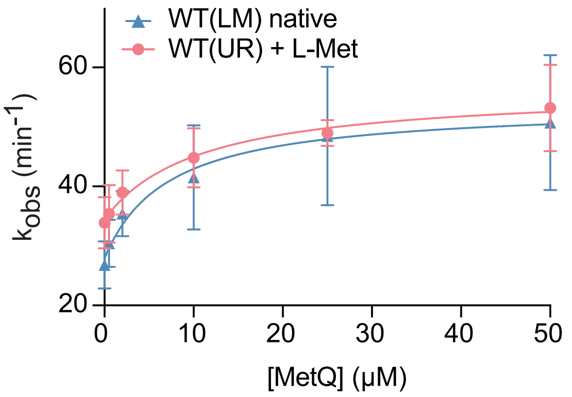


**Supporting Figure 4. MetQ stimulated ATPase activity of MetNI nanodiscs.** 32 nM N295A MetNI nanodiscs were incubated with 5 mM MgATP and varying concentrations of MetQ. Native WT MetQ containing co-purified L-Met (blue) was compared with unfolded-refolded WT MetQ with 100 μM exogenous L-Met (red). $K_m$(MetQ) values are 6 ± 2 μM and 14 ± 9 μM, respectively. $k_{cat}$(MetQ) values are 50 ± 20 min$^{-1}$ and 60 ± 10 min$^{-1}$, respectively.

### Figure S5

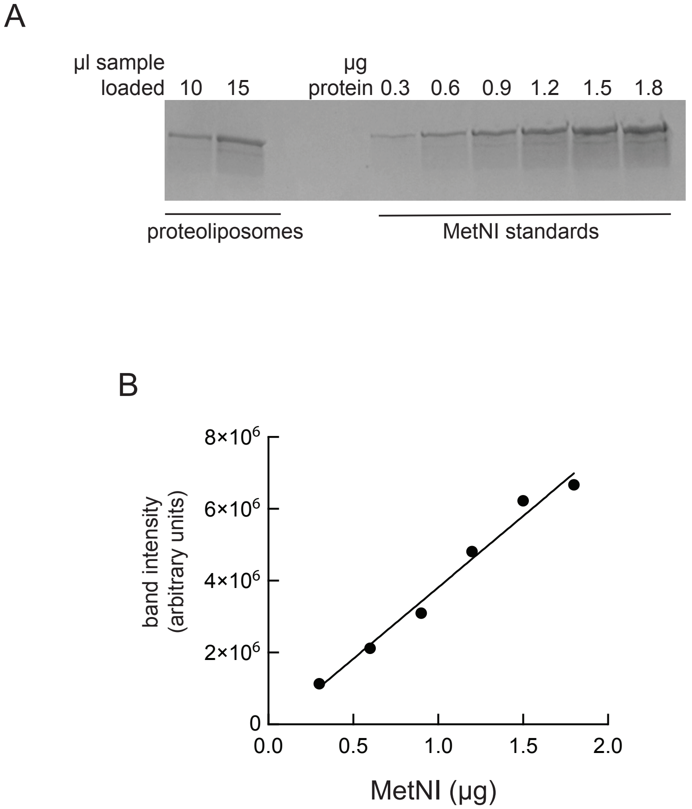


**Supporting Figure 5. Representative SDS-PAGE for liposome quantification.** (a) Coomassie-stained gel containing different amounts of a liposome preparation compared to known standards. (b) Standard curve for interpretation of data in (a). $R^2$ value is 0.9822.
